# Genetic Diversity and Genomic Epidemiology of SARS-COV-2 in Morocco

**DOI:** 10.1101/2020.06.23.165902

**Authors:** Bouabid Badaoui, Khalid Sadki, Chouhra Talbi, Salah Driss, Lina Tazi

## Abstract

COVID-A9 is an infection disease caused by severe acute respiratory syndrome coronavirus 2 (SARS-CoV-2), declared as a pandemic due to its rapid expansion worldwide. In this study we investigate the genetic diversity and genomic epidemiology of SARS-CoV-2 using 22 virus genome sequences reported by three different laboratories in Morocco till the date 07/06/2020 as well as (40366) virus genomes from all around the world.

The SARS-CoV-2 genomes from Moroccan patients revealed 62 mutations of which 30 were missense mutations. The mutations Spike_D614G and NSP12_P323L were present in all the 22 analyzed sequences, followed by N_G204R and N_R203K which occurred in 9 among the 22 sequences. The mutations NSP10_R134S, NSP15_D335N, NSP16_I169L, NSP3_L431H, NSP3_P1292L and Spike_V6F occurred one time in our sequences with no record in other sequence worldwide. These mutations should be investigated to figure out their potential effects on all around the world virulence. Phylogenetic analyses revealed that Moroccan SARS-CoV-2 genomes included 9 viruses pertaining to clade 20A, 9 to clade 20B and 2 to clade 20C. This finding suggest that the epidemic spread in Morocco did not show a predominant SARS-CoV-2 route. For multiple and unrelated introductions of SARS-CoV-2 into Morocco via different routes have occurred, giving rise to the diversity of virus genomes in the country. Furthermore, very likely, the SARS-CoV-2 virus circulated in cryptic way in Morocco starting from the fifteen January before the discovering of the first case the second of March.

## Introduction

An outbreak of respiratory illness, named COVID-19 disease, caused by SARS-CoV2 virus, have been reported in Wuhan, Hubei Province, China the beginning of December 2019 and then quickly spread to the rest of the world. On 30 January 2020, the WHO declared that the coronavirus disease constitutes a Public Health Emergency of International Concern.

This pandemic has infected more than 7,725,457 people around the world and caused more than 427,683 deaths as of June 7th, 2020 (https://www.worldometers.info/coronavirus/). This disease was confirmed to have spread to Morocco on 2 March 2020, when the first case COVID-19 case was confirmed. Till June 7, 2020 Morocco reported 8692 confirmed cases and 212 deaths. The most important feature of this disease is expressed by high-level of inflammatory response including pro-inflammatory cytokines especially in severe form which cause pneumonia and severe acute respiratory syndrome (Andersen et al. 2020). SARS-CoV-2 genome has a size of 29.8-29.9 kb harboring a long 5’ end that contains orf1ab. The 3′ end codes for the structural proteins: small envelope (E) protein, matrix (M) protein, spike (S) protein and nucleocapsid (N) protein. Furthermore, the SARS-CoV-2 contains six accessory proteins, encoded by ORF3a, ORF6, ORF7a, ORF7b, and ORF8 genes (Khailany, Safdar, and Ozaslan 2020). Compared to SARS-CoV, SARS-CoV-2 has high transmission and less pathogenicity (Fu, Cheng, and Wu 2020) for, hitherto, unknown reasons.

Viruses evolution in nature comes about through manifold mechanisms of which nucleotide substitution is of uttermost importance (Lauring and Andino 2010). Furthermore, genomic epidemiology of emerging viruses is of chief relevance for capturing virus evolution and spread (Fraser et al. 2009, and gardy et al. 2018). This approach has been proved to be very efficient during the Ebola virus disease epidemic in West Africa (Baize et al. 2014) as well as Zika virus in Brazil (Faria et al. 2017).

To investigate the mutations underlying the evolution of SARS-CoV-2 in Morocco, 40390 genomes sequences of SARS-CoV-2, of which 22 were from Moroccan patients, were collected from GISAID (https://www.gisaid.org/) and used to study the diversity and genomic epidemiology of SARS-CoV-2 in Morocco.

## Material and Methods

### SARS-CoV-2 diversity and phylogenetic

To assess the genetic variation of SARS-CoV-2 in Morocco, a total of 40390 complete genomes of SARS-CoV-2 as well their corresponding metadata were retrieved from GISAID database (Shu and McCauley 2017) and analyzed using the Nextstrain bioinformatics pipeline (Hadfield et al. 2018):

The platform Augur was used to perform a multi-alignment using all the genomes via MAFFT (Katoh et al. 2002) and (‘Wuhan-Hu-1/2019’, ‘Wuhan/WH01/2019’) as reference genomes. The phylogeny was built by maximum likelihood using IQTREE (Nguyen et al. 2015). For comparative analyze of SARS-Cov-2 genome sequences we have used the following protocol:

i) Collection of SARS-CoV-2 Sequences and the corresponding metadata from the GISAID database. ii) Filtering the SARS-CoV-2 Sequences to exclude inaccurate sequences based on missing bases and sequence length and to set a fixed number of samples per group according to their similarities. In this stage, 20 sequences from the 22 corresponding to the Moroccan patients remained for the subsequent analyses. iii) Performing a multi sequence alignment via MAFFT. vi) Inferring a phylogenetic tree from the multi-sequence alignment, getting a time-resolved tree via TreeTime and inferring ancestral traits sequences v) Identifying the mutations.

## Results

### Genetic diversity and evolution of SARS-CoV-2 in Morocco

In this study, the analysis of 20 genomes from Moroccan patients revealed 62 mutations (Supplementary Table 1) of which 30 were missense mutations (Supplementary Table 2). Among those mutations, Spike_D614G and NSP12_P323L were present in all the 22 analyzed sequences, followed by N_G204R and N_R203K which occurred in 9 sequences among the 22 (Table 1).

Till the date of the 7th June 2020, the mutations Spike_D614G and NSP12 P323L already occurred 23612 and 23543 times, respectively (70.93% and 70.90% worldwide, respectively) in 75 countries. Furthermore, the Mutations N R203K and N G204R already occurred 8744 and 8715 times, respectively (26.31% and 8715 of worldwide, respectively) in 65 countries https://www.gisaid.org/.

The mutation Spike M1237I found one time in our sequences already occurred 11 times worldwide (0.03% of all samples with Spike sequence) in six countries.

Other mutations NSP12_M196I, NSP3_A1819V, M_L13F, NSP14_D324A, NSP14_T75I and NSP5_V125I occurred one time in our sequences and have been reported in only one sequence in the GISAID database (40368 genome sequences). Therefore, NSP12_M196I, NSP3_A1819V, M_L13F, NSP14_D324A, NSP14_T75I and NSP5_V125I occurred one time in India (hCoV-19/India/GBRC46/2020); Poland (hCoV-19/Poland/PL_P15/2020); Switzerland/100144/2020; United Kingdom / England EPI_ISL_423899, Australia: EPI_ISL_426742 and South America: EPI_ISL_445334, respectively.

The mutations NSP10_R134S, NSP15_D335N, NSP16_I169L, NSP3_L431H, NSP3_P1292L and Spike_V6F occurred one time in our sequences with no record in other sequences all around the world.

### Genomic epidemiology of SARS-CoV-2 in Morocco

Herein we report on the putative history of SARS-CoV-2 transmission in Morocco as revealed by genomic epidemiology (Figure 1 and 2). The virus starts to circulate in Morocco, around the beginning of February more likely from Belgium, Spain and France (Clade 20C was the first one to be introduced to Morocco); around the 04/03/2020, new infected cases came from Belgium to Morocco (Clade 20A), around the 12/03/2020, other infected cases entered Morocco coming from France and passing through Spain (Clade 20A). Three days later (14/03/2020) infected cases reached Morocco from Switzerland (Clade A). Around the 17/03/2021, infected cases reached Morocco from USA (Clade 20A and 20B) and Germany (Clade 20B). Around the 22/03/2020 infected cases reached Morocco from Taiwan (Clade A). Other infected arrivals have been registered between the 16th and 19th April 2020 from USA and Vietnam, more likely through sea trades that continued during the lockdown.

**Figure 1.**
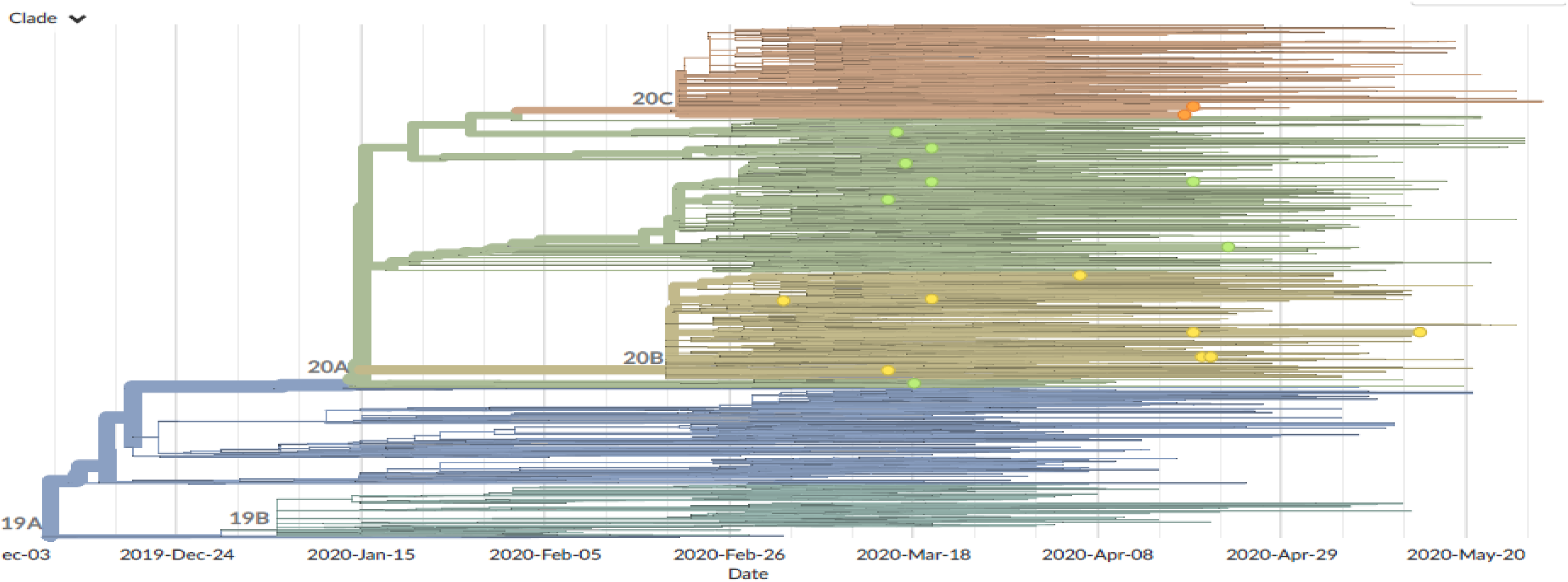
Phylogeny of 40390 SARS-CoV-2 viruses collected from GISAID database. The viruses collected from Morocco are highlighted, in colorful dots, according to the clades to which they pertain. Clustering of related viruses indicates community transmission after an introduction event. Morocco witnessed three major transmission events

**Figure 2.**
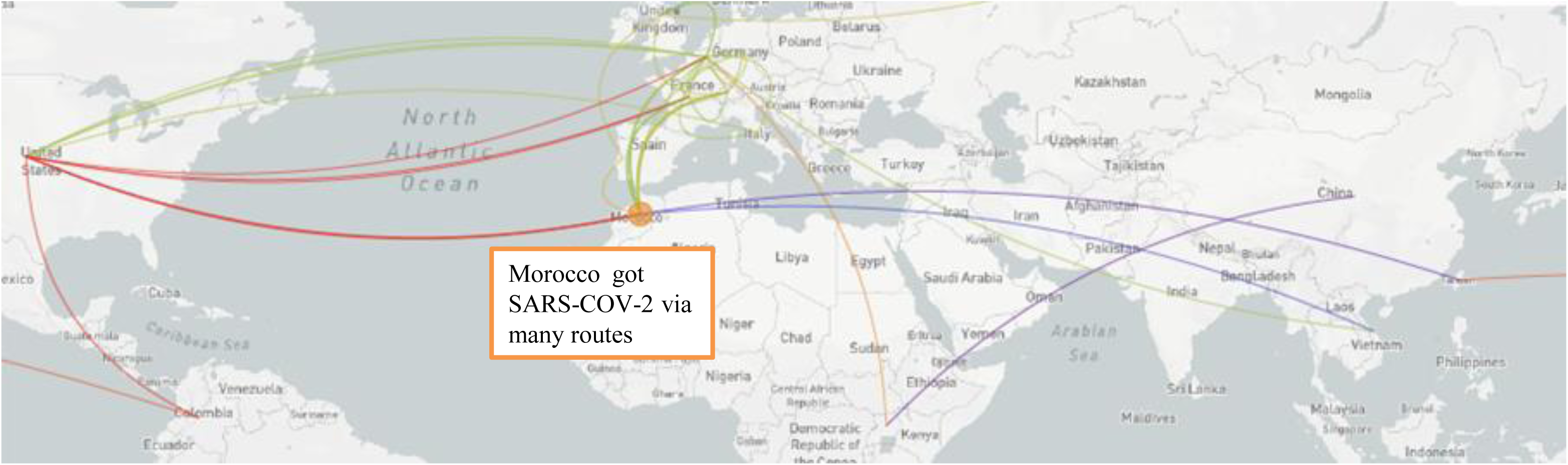
Narrative of SARS-COV-2 transmission in Morocco according the genomic epidemiology approach highlighted by the phylogeny of 40390 SARS-CoV-2 viruses collected from GISAID database.

The Moroccan genomes analyzed in this study possessed between 4 and 16 mutations relative to the common ancestor [Wuhan-Hu-1/2019’, ‘Wuhan/WH01/2019’] (Figure 3). This pattern got a molecular evolution rate of 25.331 substitution per year which is consistent with what was reported before (Rambaut 2020).

**Figure 3.**
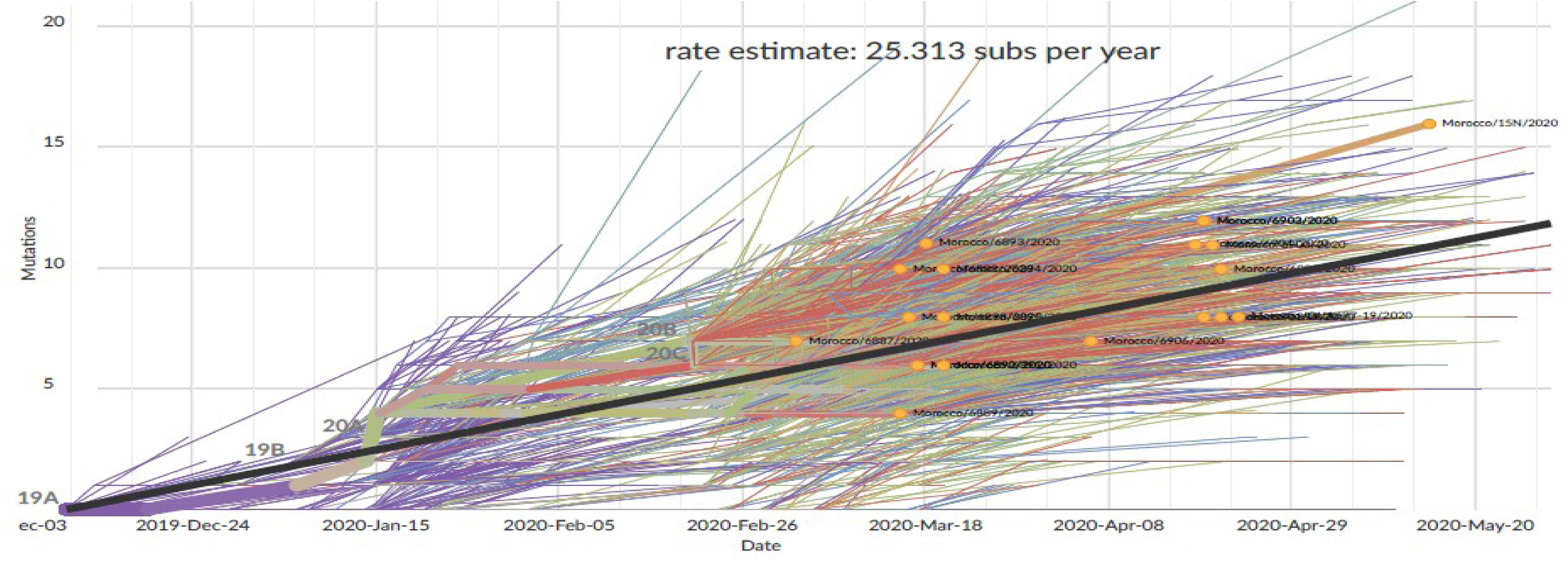
Clock Diagram showing the evolution of mutations number in SARS-COV-2 strains. The viruses collected in Morocco, colored in orange, got between 4 and 16 mutations compared to the reference sequence.

Phylogenetic analysis (Figure 1) shows that the SARS-CoV-2 genomes from Moroccan patients generated in this study are dispersed across the evolutionary tree of SARS-CoV-2 viruses, estimated from 40390 genomes available on GISAID as of June 07th, 2020. These viruses included 9 viruses from clade 20A, 9 from Clade 20B and 2 viruses from clade 20C (Figure 4). This shows that the epidemic spread in Morocco do not show a predominant SARS-CoV-2 lineage. However, it’s more likely that the virus circulated in hidden way around the beginning of February before the discovering of the first case the 2^th^ of March.

**Figure 4.**
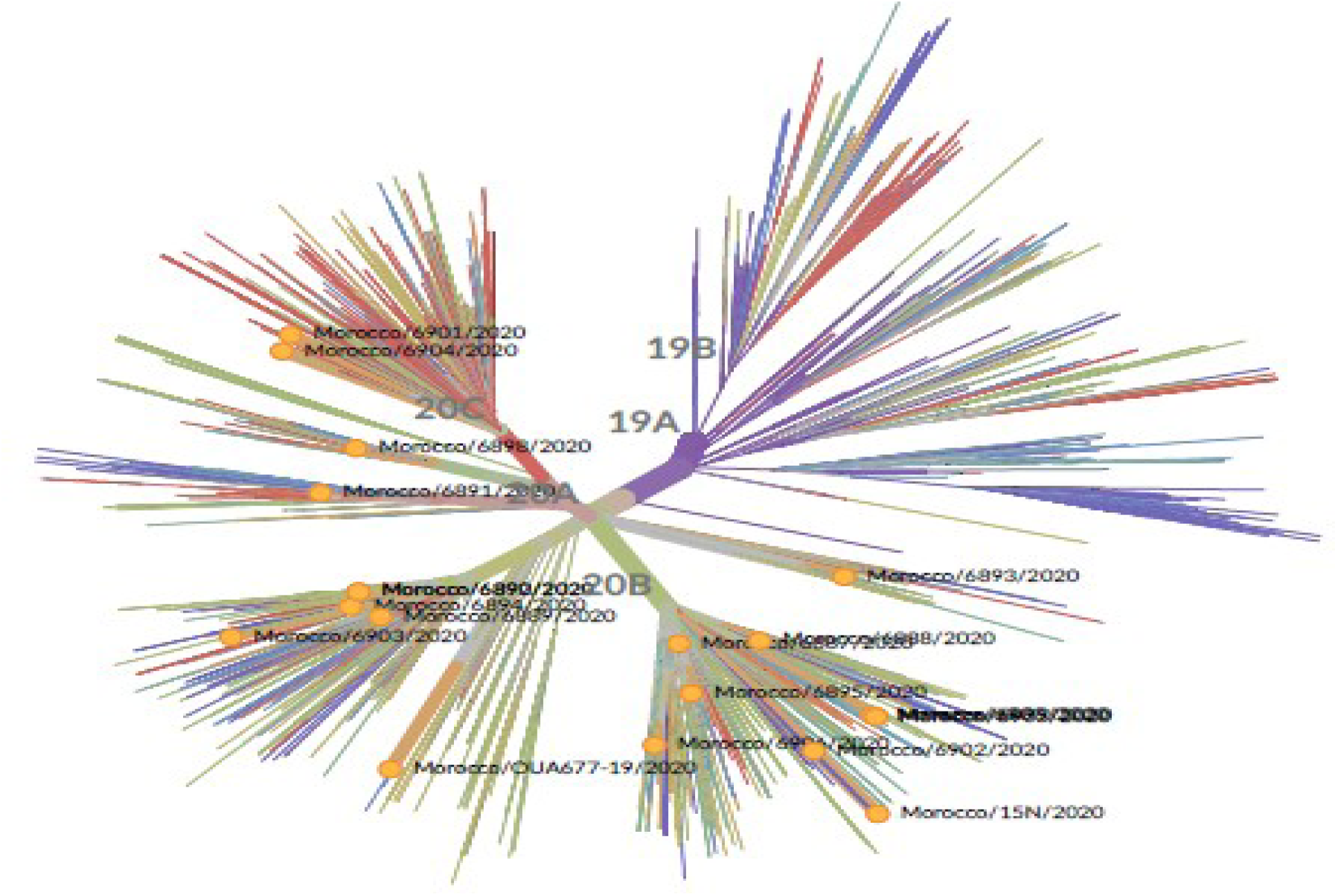
Radial tree for the Phylogeny of 40390 SARS-CoV-2 viruses collected from GISAID database. The viruses collected from Morocco are highlighted, in orange dots, pertain to three clades 20A,20B and 20C.

## Discussion

Among the missense mutations, Spike_D614G and NSP12 P323L, N R203K and N G204R occurred with high frequency worldwide. This, verisimilarly, contributes to increase the SARS-CoV-2 transmissibility. In fact, Throughout its evolution within the host, the virus seeks to proliferate efficiently whilst concurrently circumventing host morbidity to set out a maximum transmission (Alizon et al. 2009). This is in concordance with the concomitant reduced pathogenicity that accompanies its transmission increase. The Spike_D614G mutation manifested a major effect on the virus efficiency to infect hosts (Junior et al. 2020 [preprint]) and showed a high ability to hinder the immune systems of hosts that already dealt with version of SARS-CoV-2 without the Spike_D614G mutation (Zhang et al. 2020 [preprint]). This aspect should be emphasized for future vaccine researches. The mutation P323L, although provoked the substitution of proline that plays a prominent role in protein folding, and aggregation, neither increased the SARS-CoV2’s infectivity nor its fitness regarding natural selection (Maitra et al 2020). This might be because the change from proline to leucine amino-acid (P323L) did not change the protein function as both amino-acids pertain to the nonpolar aliphatic R groups.

Other missense mutations occurred either with small frequency like Spike M1237I that was reported 11 times all over the world or have been reported only one time worldwide like NSP12_M196I, NSP3_A1819V, M_L13F, NSP14_D324A, NSP14_T75I and NSP5_V125I. These mutations seems to be rare and may represent the adaptation expression of the virus in front of specific genetic backgrounds of the host, climate conditions or other unknown factors. In what concerns the mutation Spike M1237I, It’s noteworthy to mention that the spike surface glycoprotein is crucial for the virus binding to receptors on the host cell (Fung and Liu 2019). This glycoprotein is also the major target of neutralizing antibodies (Chen et al. 2020). Hence, mutations in the spike surface glycoprotein could provoke change in SARS-CoV-2 antigenicity.

The mutations NSP10_R134S, NSP15_D335N, NSP16_I169L, NSP3_L431H, NSP3_P1292L and Spike_V6F that occurred in the Moroccan sequences with no record in other sequences worldwide, should get a careful attention and should be investigated to figure out their potential effects on the SARS-CoV-2 virulence as well as their association with immunological and clinical symptoms. Careful attention should be afforded to the mutation Spike_V6F as the structural protein spike might essential for the virus ability to infect the hosts and was used as an important target for vaccine development (Ahmed, Quadeer, and McKay 2020).

Genomic epidemiology, using Nextstrain, applied to SARS-CoV-2 transmission in Morocco revealed many aspects of the epidemic that was already known by the official authorities like the introduction of SARS-CoV-2 strains to Morocco from Belgium, Spain and France at the beginning of the epidemic. However, this analysis highlighted many other events hitherto remained unknown like the arrival of strains from USA and Vietnam after the lockdown, possibly, through sea trades.

SARS-CoV-2 genomes from Moroccan patients are dispersed across the evolutionary tree of SARS-CoV-2 viruses with 9 viruses pertaining to clade 20A, 9 to Clade 20B and 2 viruses to clade 20C. This suggests that multiple and unrelated introductions of COVID-19 into Morocco via different routes have occurred, giving rise to the diversity of virus lineages reported in this study. This finding suggest that different SARS-CoV-2 strains, with different mutation patterns, coexist in Morocco. The contribution of each of those mutations needs to be investigated in order to figure out possible drug resistance and eventually different SARS-CoV-2 mortality rates related to those mutations.

It’s tempting to speculate that the specific evolutionary profiles of the SARS-COV-2 in Morocco might be the product of the interaction between its evolutionary routes before reaching the country and its adaptation to the Moroccan genetic background.

This genomic survey of SARS-CoV-2 in Morocco has at least two limitations. First, the sampling size is relatively weak which might hinder a compelling interpretation of the epidemiological situation in the country. Second, all the samples analyzed, except one, were obtained from public health laboratories and thus may not be representative of the general population.

## Conclusions

The genetic analysis of the SARS-CoV-2 genomes from Moroccan patients revealed some new mutations with no aforementioned record in other sequences worldwide. These mutations should be investigated to figure out their potential effects on the hCoV19 virulence and to evaluate their impacts on the immune response. The Phylogenetic analyses revealed that the COVID-19 spread in occurred through multiple and unrelated introductions of COVID-19 into Morocco via different routes. SARS-CoV-2 virus might have circulated in cryptic way in Morocco around the beginning of February before the discovering of the first case the 2^d^ of March.

