## Supplementary Tables for "Genetic Diversity and Genomic Epidemiology of SARS-COV-2 in Morocco"

Table S1. SARS-CoV-2 genomes from Moroccan patients used in the current study

| Query | Strand | %N | Length(nt) | Length (aa) | #Muts | %Muts | #Unique Muts | %UniqueMuts | #ExistingMuts | %Existing Muts |
|---|---|---|---|---|---|---|---|---|---|---|
| **hCoV-19/Morocco/6887/2020\|EPI_ISL_459965\|2020-03-03** | positive | 0.01 % | 29903 | 9710 | 4 | 0.04 % | 0 | 0.00% | 4 | 0.04% |
| **hCoV-19/Morocco/6888/2020\|EPI_ISL_459966\|2020-03-15** | positive | 0.00 % | 29903 | 9710 | 7 | 0.07 % | 1 | 0.01% | 6 | 0.06% |
| **hCoV-19/Morocco/6889/2020\|EPI_ISL_459967\|2020-03-15** | positive | 0.90 % | 29903 | 9710 | 2 | 0.02 % | 0 | 0.00% | 2 | 0.02% |
| **hCoV-19/Morocco/6890/2020\|EPI_ISL_459968\|2020-03-17** | positive | 0.72 % | 29903 | 9710 | 2 | 0.02 % | 0 | 0.00% | 2 | 0.02% |
| **hCoV-19/Morocco/6891/2020\|EPI_ISL_459969\|2020-03-20** | positive | 1.20 % | 29903 | 9109 | 3 | 0.03 % | 0 | 0.00% | 3 | 0.03% |
| **hCoV-19/Morocco/6892/2020\|EPI_ISL_459970\|2020-03-17** | positive | 1.24 % | 29903 | 9710 | 3 | 0.03 % | 1 | 0.01% | 2 | 0.02% |
| **hCoV-19/Morocco/6893/2020\|EPI_ISL_459971\|2020-03-18** | positive | 1.39 % | 29903 | 9581 | 7 | 0.07 % | 3 | 0.03% | 4 | 0.04% |
| **hCoV-19/Morocco/6894/2020\|EPI_ISL_459972\|2020-03-20** | positive | 0.00 % | 29903 | 9710 | 4 | 0.04 % | 0 | 0.00% | 4 | 0.04% |
| **hCoV-19/Morocco/6895/2020\|EPI_ISL_459973\|2020-03-20** | positive | 0.00 % | 29903 | 9710 | 5 | 0.05 % | 1 | 0.01% | 4 | 0.04% |
| **hCoV-19/Morocco/6896/2020\|EPI_ISL_459974\|2020-03-20** | positive | 0.06 % | 29903 | 9710 | 4 | 0.04 % | 0 | 0.00% | 4 | 0.04% |
| **hCoV-19/Morocco/6897/2020\|EPI_ISL_459975\|2020-03-21** | positive | 0.08 % | 29903 | 9710 | 3 | 0.03 % | 1 | 0.01% | 2 | 0.02% |
| **hCoV-19/Morocco/6898/2020\|EPI_ISL_459976\|2020-03-16** | positive | 0.02 % | 29903 | 9710 | 4 | 0.04 % | 0 | 0.00% | 4 | 0.04% |
| **hCoV-19/Morocco/6899/2020\|EPI_ISL_459977\|2020-04-21** | positive | 0.00 % | 29903 | 9710 | 5 | 0.05 % | 0 | 0.00% | 5 | 0.05% |
| **hCoV-19/Morocco/6900/2020\|EPI_ISL_459978\|2020-04-20** | positive | 0.04 % | 29903 | 9710 | 7 | 0.07 % | 1 | 0.01% | 6 | 0.06% |
| **hCoV-19/Morocco/6901/2020\|EPI_ISL_459979\|2020-04-19** | positive | 0.00 % | 29903 | 9710 | 4 | 0.04 % | 0 | 0.00% | 4 | 0.04% |
| **hCoV-19/Morocco/6902/2020\|EPI_ISL_459980\|2020-04-19** | positive | 0.00 % | 29903 | 9710 | 6 | 0.06 % | 1 | 0.01% | 5 | 0.05% |
| **hCoV-19/Morocco/6903/2020\|EPI_ISL_459981\|2020-04-19** | positive | 0.00 % | 29903 | 9710 | 5 | 0.05 % | 1 | 0.01% | 4 | 0.04% |
| **hCoV-19/Morocco/6904/2020\|EPI_ISL_459982\|2020-04-18** | positive | 0.02 % | 29903 | 9710 | 6 | 0.06 % | 2 | 0.02% | 4 | 0.04% |
| **hCoV-19/Morocco/6905/2020\|EPI_ISL_459983\|2020-04-21** | positive | 0.05 % | 29903 | 9710 | 6 | 0.06 % | 1 | 0.01% | 5 | 0.05% |
| **hCoV-19/Morocco/6906/2020\|EPI_ISL_459984\|2020-04-06** | positive | 0.00 % | 29903 | 9710 | 4 | 0.04 % | 0 | 0.00% | 4 | 0.04% |
| **hCoV-19/Morocco/OUA677-19/2020\|EPI_ISL_451400\|2020-04-23** | positive | 0.00 % | 29934 | 9710 | 4 | 0.04 % | 0 | 0.00% | 4 | 0.04% |
| **hCoV-19/Morocco/15N/2020\|EPI_ISL_458150\|2020-05-15** | positive | 0.13 % | 29838 | 9710 | 7 | 0.07 % | 3 | 0.03% | 4 | 0.04% |

Table S2 . Nucleotide mutations in SARS-CoV-2 from Moroccan patients

| strain | divergence | excess divergence | #Ns | #gaps | clusters | Gaps | SNPs |
|---|---|---|---|---|---|---|---|
| **Morocco/ 15N/2020** | 16 | 4.63 | 38 | 0 | | | 240,312,690,3036,3873,4010,5496,8174,12099,14407,21512,23402,26558 |
| **Morocco/ 6903/2020** | 12 | 2.41 | 17 | 0 | | | 240,1683,3036,4233,6403,8207,14407,15323,19009,22362,23402,29361 |
| **Morocco/ 6902/2020** | 12 | 2.41 | 17 | 0 | | | 240,312,690,3036,8174,14407,16607,23402,25272,28880,28881,28882 |
| **Morocco/ 6900/2020** | 11 | 1.34 | 0 | 0 | | | 221,240,3036,3372,14027,14407,19812,23402,28880,28881,28882 |
| **Morocco/ 6893/2020** | 11 | 3.6 | 433 | 0 | | | 240,3036,10522,13423,14096,14407,14579,19297,21162,21165,23402 |
| **Morocco/ 6904/2020** | 11 | 1.48 | 0 | 0 | | | 240,1058,3036,9840,9966,14407,20622,21577,23402,25562,27166 |
| **Morocco/ 6894/2020** | 10 | 2.47 | 0 | 0 | | | 240,1683,3036,6403,8207,14407,15323,22362,23402,29361 |
| **Morocco/ 6896/2020** | 10 | 2.47 | 0 | 0 | | | 240,1683,3036,6403,8207,14407,15323,22362,23402,29361 |
| **Morocco/ 6888/2020** | 10 | 2.81 | 17 | 0 | | | 240,3036,10426,14407,14785,20388,23402,28880,28881,28882 |
| **Morocco/ 6905/2020** | 10 | 0.27 | 2 | 0 | | | 221,240,3036,3372,14027,14407,23402,28880,28881,28882 |
| **Morocco/ 6895/2020** | 8 | 0.47 | 17 | 0 | | | 240,3036,14407,18262,23402,28880,28881,28882 |
| **Morocco/ 6899/2020** | 8 | -1.73 | 17 | 0 | | | 240,3036,3372,14407,23402,28880,28881,28882 |
| **Morocco/ 6901/2020** | 8 | -1.59 | 17 | 0 | | | 202,240,1058,1665,3036,14407,23402,25562, |
| **Morocco/ 6898/2020** | 8 | 0.74 | 0 | 0 | | | 240,2415,3036,14407,16205,23402,25562,29772 |
| **Morocco/ OUA677-19/2020** | 8 | -1.86 | 0 | 0 | | | 197,240,3036,14407,15379,18507,23402,28737 |
| **Morocco/ 6887/2020** | 7 | 0.63 | 12 | 0 | | | 240,3036,14407,23402,28880,28881,28882 |
| **Morocco/ 6906/2020** | 7 | -1.7 | 17 | 0 | | | 240,3036,14407,23402,28880,28881,28882 |
| **Morocco/ 6897/2020** | 6 | -1.6 | 0 | 1 | | 29724-29725 | 240,3036,6593,14407,20267,23402 |
| **Morocco/ 6890/2020** | 6 | -1.33 | 204 | 0 | | | 240,3036,3240,14407,15323,23402 |
| **Morocco/ 6891/2020** | 6 | -1.53 | 331 | 0 | | | 240,3036,14407,18876,23402,25562 |
| **Morocco/ 6892/2020** | 6 | -1.33 | 350 | 0 | | | 240,3036,3240,14407,15323,23402 |
| **Morocco/ 6889/2020** | 4 | -3.19 | 289 | 0 | | | 240,3036,14407,23402 |

| | | | | | | | |
|---|---|---|---|---|---|---|---|
| **Wuhan/ WH01/2019** | 2 | 0.29 | 0 | 0 | | | 6967,11763 |
| **Wuhan-Hu-1/2019** | 0 | -1.71 | 0 | 0 | | | 240,312,690,3036,3873,4010,5496,8174,12099,14407,21512,23402,26558 |

Table S3 . Non synonymous mutations in SARS-CoV-2 from Moroccan patients

| Mutation | Count |
|---|---|
| NSP12_P323L | 22 |
| Spike_D614G | 22 |
| N_G204R | 9 |
| N_R203K | 9 |
| NS3_Q57H | 4 |
| NSP3_D218E | 3 |
| NSP3_T1830I | 3 |
| NSP3_V1229F | 3 |
| NSP12_M196I | 2 |
| NSP2_T85I | 2 |
| NSP3_A1819V | 2 |
| M_L13F | 1 |
| NSP10_R134S | 1 |
| NSP12_A449V | 1 |
| NSP12_E922D | 1 |
| NSP12_M380I | 1 |
| NSP12_S647I | 1 |
| NSP14_D324A | 1 |
| NSP14_L157F | 1 |
| NSP14_T75I | 1 |
| NSP14_Y420C | 1 |
| NSP15_D335N | 1 |
| NSP15_P65S | 1 |
| NSP15_R257C | 1 |
| NSP16_I169L | 1 |
| NSP3_L1417J | 1 |
| NSP3_L431H | 1 |
| NSP3_P1292L | 1 |
| NSP5_V125I | 1 |
| NSP5_V157L | 1 |
| Spike_M1237I | 1 |
| Spike_V6F | 1 |
